# Optimal synaptic dynamics for memory maintenance in the presence of noise

**DOI:** 10.1101/2020.08.19.257220

**Authors:** Dhruva V Raman, Timothy O’Leary

## Abstract

Synaptic connections in many brain areas have been found to fluctuate significantly, with substantial turnover and remodelling occurring over hours to days. Remarkably, this flux in connectivity persists in the absence of overt learning or behavioural change. What proportion of these ongoing fluctuations can be attributed to systematic plasticity processes that maintain memories and neural circuit function? We show under general conditions that the optimal magnitude of systematic plasticity is typically less than the magnitude of perturbations due to internal biological noise. Thus, for any given amount of unavoidable noise, 50% or more of total synaptic turnover should be effectively random for optimal memory maintenance. Our analysis does not depend on specific neural circuit architectures or plasticity mechanisms and predicts previously unexplained experimental measurements of the activity-dependent component of ongoing plasticity.

## INTRODUCTION

Learning depends systematic changes to the connectivity and strengths of synapses in neural circuits. This has been shown across experimental systems (Moczulska et al., 2013; Lai et al., 2012; Hayashi-Takagi et al., 2015) and is assumed by most theories of learning (Hebb, 1949; Bienenstock et al., 1982; Gerstner et al., 1996).

Neural circuits are required not only to learn, but to retain previously learned information. One might therefore expect synaptic stability in the absence of an explicit learning signal. However, many recent experiments in multiple brain areas have documented substantial ongoing synaptic modification in the absence of any obvious learning or change in behaviour (Attardo et al., 2015; Pfeiffer et al., 2018; Holtmaat et al., 2005; Loewenstein et al., 2015; Yasumatsu et al., 2008; Loewenstein et al., 2011).

It is natural to ask whether this apparently irreducible flux in neural connectivity is due to random biochemical noise, or due to systematic plasticity processes that have not been accounted for. A number of experimental studies have attempted to dissect and quantify both systematic and random synaptic changes at the level of synaptic physiology, either by directly interfering with synaptic plasticity or by correlating changes to circuit-wide measurements of ongoing physiological activity (Nagaoka et al., 2016; Quinn et al., 2019; Yasumatsu et al., 2008; Minerbi et al., 2009; Dvorkin and Ziv, 2016). Consistently, these studies find that the total rate of ongoing synaptic change is reduced by only 50% or less in the absence of neural activity. Similarly, systematic processes that are correlated across synapses only account for less than 50% of ongoing changes. Thus the bulk of ongoing synaptic fluctuations seem to be due to internal biological noise.

Put another way, these experimental findings imply that at steady state, systematic plasticity processes exert a weaker effect on synaptic strength than random fluctuations. This is surprising, because maintenance of neural circuit properties and learned behaviour would intuitively require random fluctuations to be dominated by systematic plasticity. To our knowledge, there is no theoretical account or model prediction that explains these observations.

In this study we consider neural circuits attempting to optimally retain previously learned information through some active, systematic plasticity process, in the presence of unavoidable, learning-independent, synaptic fluctuations. We conduct a first-principles mathematical analysis that is independent of specific plasticity mechanism and circuit architectures. We find that the magnitude of systematic plasticity should not exceed those of the intrinsic fluctuations, in direct agreement with experimental data. Furthermore, these fluctuations should dominate when systematic plasticity mechanisms are relatively precise, suggesting that random fluctuations will often dominate synaptic dynamics in neural circuits that exhibit learning-related plasticity. We validate these theoretical predictions in simulations. Together, our results provide a simple and general theory that explains a number of convergent but puzzling experimental findings, and suggest that synaptic plasticity mechanisms are optimised for the dynamic maintenance of stored information.

## RESULTS

We begin with a brief survey of quantitative, experimental measurements of synaptic dynamics. We focused on studies that measured ‘baseline’ synaptic changes that occur outside of any behavioural learning paradigm, and which controlled for stimuli that induce widespread adaptive changes in synaptic strength.

Table 1 shows a breakdown of measured baseline synaptic modifications from multiple studies and brain preparations. We note that there is large heterogeneity in the rates of baseline synaptic turnover across preparations and experimental conditions. For example, the expected lifetime of synapses in adult mouse hippocampus was estimated as 1 – 2 weeks Attardo et al. (2015); Pfeiffer et al. (2018), while > 70% of synapses in mouse barrel cortex persisted over 18 months Zuo et al. (2005).

**Table 1.** Synaptic plasticity rates across experimental models, and the effect of activity suppression

| Reference | Experimental system | Total baseline synaptic change | % synaptic change that is random / learning-independent |
| --- | --- | --- | --- |
| Pfeiffer et al. (2018) | Adult mouse hippocampus | 40% turnover over 4 days | NA |
| Loewenstein et al. (2011) | Adult mouse auditory cortex | > 70% of spines changed size by > 50% over 20 days | NA |
| Zuo et al. (2005) | Adult mouse (barrel, primary motor, frontal) cortex | 3 – 5% turnover over 2 weeks for all regions. $73.9 \pm 2.8\%$ of spines stable over 18 months (barrel cortex) | NA |
| Nagaoka et al. (2016) | Adult mouse visual cortex | 8% turnover per 2 days in visually deprived environment. 15% in visually enriched environment. 7 – 8% in both environments under pharmacological suppression of spiking. | $\approx 50\%$ (turnover) |
| Quinn et al. (2019) | Glutamatergic synapses, dissociated rat hippocampal culture | $28.2 \pm 3.7\%$ of synapses formed over 24 hour period. $28.6 \pm 2.3\%$ eliminated. Activity suppression through tetanus neurotoxin -light chain. Plasticity rate unmeasured. | $\approx 75\%$ (turnover) |
| Yasumatsu et al. (2008) | CA1 pyramidal neurons, primary culture, rat hippocampus | Measured rates of synaptic turnover and spine-head volume change. Baseline conditions vs activity suppression (NMDAR inhibitors). Turnover rates: $32.8 \pm 3.7\%$ generation/elimination per day (control) vs $22.0 \pm 3.6\%$ (NMDAR inhibitor). Rate of spine-head volume change: | $\approx 67 \pm 17\%$ (turnover). Size-dependent, but consistently > 50% (spine-head volume) |
| Dvorkin and Ziv (2016) | Glutamatergic synapses in cultured networks of mouse cortical neurons | Partitioned commonly innervated (CI) synapses sharing same axon and dendrite, and non-CI synapses. Quantified covariance in fluorescence change for CI vs non-CI synapses to estimate relative contribution of activity histories to synaptic remodelling | 62 – 64% (plasticity) |
| Minerbi et al. (2009) | Rat cortical neurons in primary culture | Created “relative synaptic remodeling measure” (RRM) based on frequency of changes in the rank ordering of synapses by fluorescence. Compared baseline RRM to when neural activity was suppressed by tetrodotoxin (TTX). RRM: 0.4 (control) vs 0.3 (TTX) after 30 hours. | $\approx 75\%$ (plasticity) |
| Ziv and Brenner (2018) | Literature review across multiple systems | “Collectively these findings suggest that the contributions of spontaneous processes and specific activity histories to synaptic remodeling are of similar magnitudes” | $\approx 50\%$ |

It is reasonable to assume that some component of the total ongoing synaptic changes measured in these experiments arise from unavoidable, noisy fluctuations. Left unchecked, such random perturbations would eventually disrupt circuit function and erase any memories stored in the synaptic weight distribution. This suggests that there should be an additional, systematic component of ongoing plasticity that compensate for the deleterious effect of such fluctuations.

In order to isolate and quantify these components, several experiments in Table 1 blocked known synaptic plasticity pathways, reduced environmental stimuli or statistically factored out the effect of ongoing neural activity. Intriguingly, the rates of ongoing synaptic change remained high. Moreover, the relative reduction in synapse dynamics was remarkably consistent: across multiple brain regions, *in vivo* and *in vitro*, and despite large methodological differences, these studies consistently reported that at least half of ongoing synaptic change persisted.

These surprising observations motivated the central question that we address in this study:

> *how much systematic plasticity is expected in a neural circuit that needs to maintain overall function on a previously learned task while being subjected to unavoidable, task-independent synaptic fluctuations?*

We emphasize that our goal is not to explain the source of the intrinsic synaptic fluctuations, the mechanism of systematic plasticity, nor the *total* magnitude of ongoing synaptic change. Our goal is to derive a general relationship between the sources of ongoing change in synapses, and in doing so explain why random synaptic fluctuations seem to dominate.

For generality, we wanted to make minimal assumptions about the mechanisms of synaptic plasticity, as well as circuit architecture and function. The problem we are considering is outlined in Figure 1a. A neural network can perform some learned task. The level of task performance depends on the the state of various network parameters, such as synaptic connections strengths, and intrinsic neuron properties. Our results are independent of which kinds of parameters are involved, so we name them ‘synaptic weights’ for convenience. Task performance can be quantified in terms of an error function, *F*[**w**(*t*)], which depends upon the vector **w**(*t*) of synaptic weights at time *t*.

**Figure 1.**
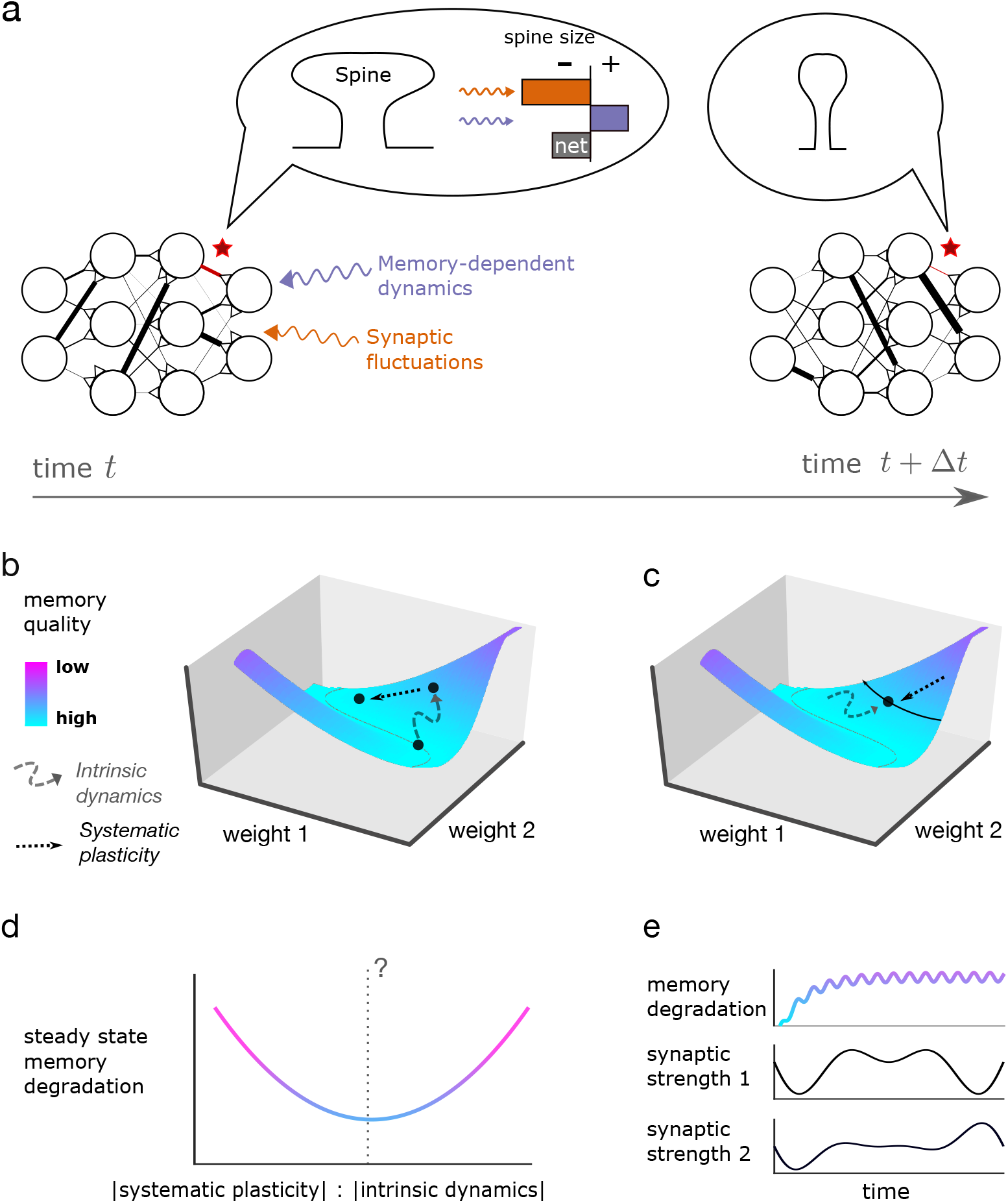
a: A learned task corresponds to an enforced input-output transformation for a neural circuit. This depends on the configuration of network parameters such as synaptic connection strengths, which change over time. Some changes arise from task-independent sources (e.g. molecular noise in the dynamic processes maintaining synapses). Unchecked, these processes will degrade task performance. Therefore some systematic ‘relearning’ term must nullify their functional effect. The combination of synaptic processes results in stable task performance, even as network parameters change systematically (e.g. see synapse labelled with red star). b: We represent task performance as a ‘landscape’. Lateral co-ordinates represent the values of network parameters (only two are visualised). Height represents task error, so lower regions correspond to better performing configurations of network parameters. Ongoing change in the network parameters corresponds to ‘wandering’ across the landscape. Near the bottom (high task performance), task-independent dynamics will tend to move upwards on the landscape. Systematic plasticity moves doenwards, since it improves task performance. c: Eventually, a steady state is reached at which the effect of the two competing terms on memory quality cancel out. The network parameters wander over a level set of the landscape. d: For a fixed magnitude of task-independent synaptic fluctuations, what magnitude of systematic plasticity maximises memory quality (i.e. gives us the ‘best level set’ on the landscape)? e: Individual synapses may experience large, systematic changes at this steady state.

We assume that at least some ongoing plasticity processes are ‘task-independent’, and collectively refer to these processes as ‘synaptic fluctuations’. Our definition of a task-independent process is one for which the probability of the process increasing or decreasing a particular synaptic weight, in a small time window, is independent of whether such a change is beneficial to task performance. One such process would be molecular noise affecting synaptic connection strengths. Another might be homeostatic mechanisms internal to each neuron. These perturb the weights in a direction *ε*[**w**(*t*)]. Our definition of ‘task-independent’ implies that such changes are on average uncorrelated with the direction of change in **w**(*t*) that would elicit maximal improvement in task performance, namely −∇*F*[**w**(*t*)]. For this reason, we will often refer to them as ‘random’ or ‘noise’ components, even though they may have a deterministic origin. By definition, we have:

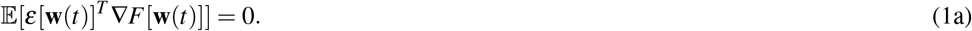

Intuitively, one would expect random fluctuations to degrade task performance. To formalise this, we will say that the network is in a **partially trained** state if a small, random change in synaptic weights satisfying equation (1a), degrades memory quality in expectation. Mathematically (see SI section 1.2), this is equivalent to the following condition:

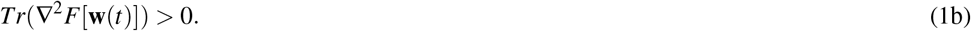

The intuition for (1b) is as follows. We conceptualise *F* as a landscape, where **w** are the coordinates of a point on the landscape, and *F*[**w**] is the height of the landscape at **w** (Figure 1b). Improving task performance corresponds to moving downhill on the landscape. Equation (1b) says a randomly chosen direction on the landscape exhibits upward curvature on average. This is because each eigenvector of ∇^2^*F*[**w**] corresponds to a direction on the landscape with curvature specified by the associated eigenvalue. If the sum of eigenvalues is positive (i.e. equation (1b)), then positive curvature is more prevalent than negative. For an isolated (local) minimum of the loss function, all eigenvalues are positive.

What is the relevance of curvature in determining the effect of random weight changes? A weight perturbation satisfying equation (1a) will be biased neither uphill nor downhill. We can pick a two-sided vector **v** along which to perturb **w** (Figure 2b). Moving in the downhill direction of **v** improves (decreases) *F*[**w**]. However, upward curvature diminishes the degree of improvement. On the other hand, moving in the uphill direction increases *F*[**w**], and this increase is magnified by upward curvature. Thus a random fluctuation along **v**, biased in neither the positive or negative direction of **v**, will decrease task performance (increase *F*[**w**]) in expectation. Equation (1b) follows by averaging over all possible directions **v** in a partially trained network.

**Figure 2.**
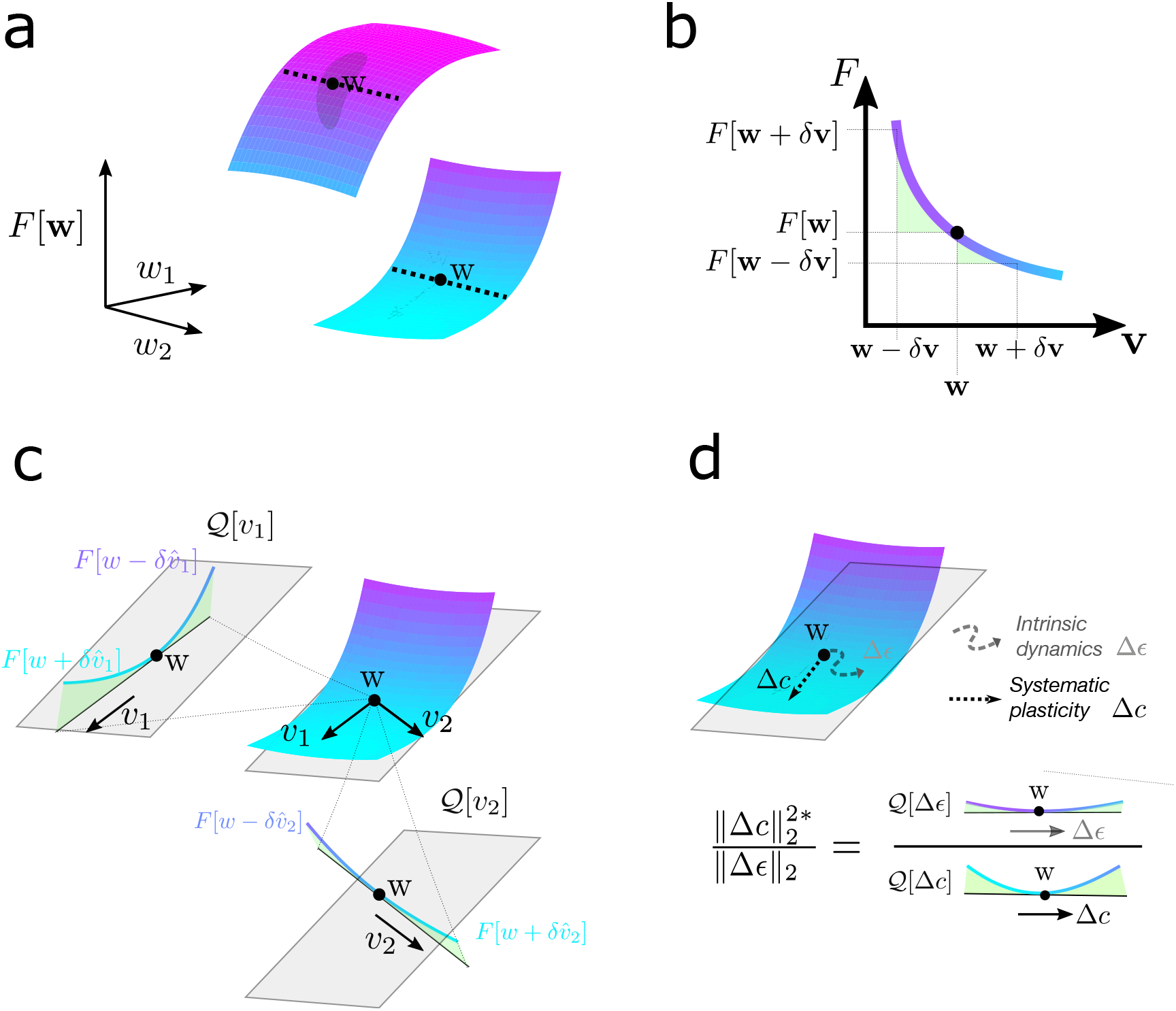
a: Two different error surfaces. Shaded patches are probability distributions of changes to **w**. The distributions are uncorrelated, in expectation, with the gradient. We see this as half the probability density lies on each side of the dotted line denoting the level set. The concave (convex) surface on top (bottom) curves downwards (upwards) in all directions, which means that a random change in **w** following the probability distribution will decrease (increase) error in expectation. If a network state is such that random changes, uncorrelated with ∇*F*[**w**], increase error in expectation, then the error surface curves upwards along the probability distribution more than it curves downwards. b: Geometrical intuition behind the operator 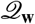. The operator charts the degree to which a direction lifts off the tangent plane (grey). In other words, the relative upward curvature of the surface in a given direction. The green, shaded areas are proportional to 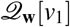, and 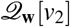. Note that the operator considers normalised directions, so does not take account the magnitude of either of these vectors. c: The weights of a networks change due to memory-independent ‘intrinsic dynamics’, as well as systematic plasticity that acts to reconsolidate the memory. What is the optimal magnitude of systematic plasticity, relative to the magnitude of the intrinsic dynamics? As 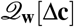 increases and/or 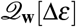 increases, the optimal magnitude of systematic plasticity increases.

This reasoning explains why synaptic fluctuations are expected to degrade task performance in a partially trained network. If such fluctuations are unavoidable but task performance is maintained, then a systematic plasticity term must counteract the fluctuations. We will denote this term **c**[**w**(*t*)]. For notational clarity, we will subsequently omit explicit dependence on **w**(*t*), i.e. **c**[**w**(*t*)] and *ε*[**w**(*t*)] become **c**(*t*) and *ε*(*t*). Overall synaptic dynamics can now be written as

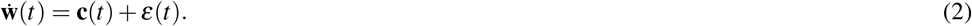

Note that many studies attempt to model the distribution of fluctuations *ε*(*t*). For instance, Statman et al. (2014); Yasumatsu et al. (2008); Loewenstein et al. (2011) identify the importance of synapse size, age, and morphology in determining fluctuation magnitude. Our study is agnostic to such considerations, as long as fluctuations are memory independent (i.e. satisfy (1a)).

The systematic plasticity term **c**(*t*) lumps together the contribution of all synaptic plasticity mechanisms that dynamically maintain the learned task. These might be referred to as ‘learning rules’ but we emphasize that most of our interest lies in the steady-state *maintenance* of a learned task, i.e. when no additional learning is occurring.

Consider the plasticity rates of **c**(*t*) and *ε*(*t*) over a small time interval, Δ*t*. We define:

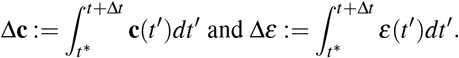

Δ**c** represents the cumulative effect of systematic plasticity and Δ*ε* the cumulative effect of fluctuations over the time interval Δ*t*. Now consider the change in memory quality during this time, Δ*F* := *F*[**w**(*t*\* + Δ*t*)] − *F*[**w**(*t*\*)]. A second order Taylor approximation of Δ*F* gives:

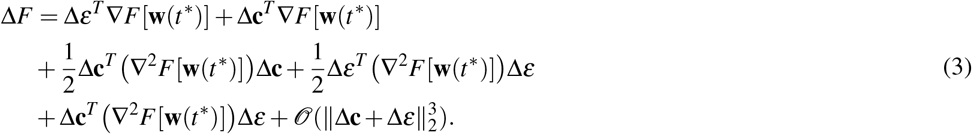

We can choose Δ*t* sufficiently small that the higher order terms in 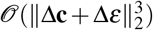 can be ignored. We do not make specific assumptions on the mechanism by which the learning rule produces synaptic changes Δ**c**. By definition, however, the effect of Δ**c** is to improve task performance. For Δ**c** to have such an effect on Δ*F*, equation (3) shows that we require

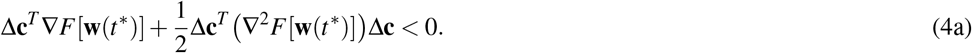

Indeed for a sufficiently small Δ*t*, we additionally require that

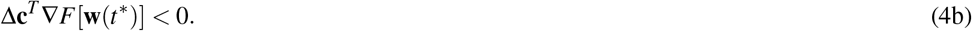

This shows that Δ**c** must be anticorrelated with the gradient ∇*F*[**w**]. Additionally, it might exploit information on ∇^2^*F*[**w**]. Geometrically, Δ**c** points in a descending direction on the loss landscape of *F*[**w**].

Our first task is to find what magnitude of systematic plasticity (i.e. ‖Δ**c**‖_2_) optimises the degree of learning Δ*F*, assuming any fixed direction 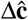. We use hats to denote normalised variables (i.e. 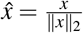). We will make extensive use of the following operator:

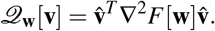

Geometrically, 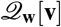 represents the relative degree of upward curvature of the loss landscape of *F* at the point **w**, in the direction **v**. This is shown graphically in Figure 2c. Note also that 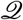 is scale invariant, i.e. 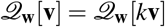 for any scalar *k* ∈ ℝ. It depends on the direction, not the magnitude, of **v**.

We can rewrite equation (3), using the operator 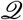 and omitting higher order terms (as previously justified):

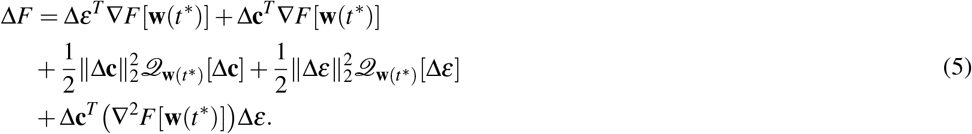

Since Δ*ε* consists of memory-independent processes, we can consider them as coming from some unknown probability distribution that is uncorrelated, in expectation, with the derivatives of *F*. Thus, any term in equation (3) that is linear in Δ*ε* disappears in expectation. In particular,

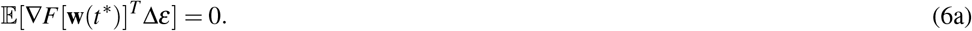

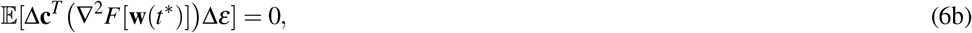

which collectively imply

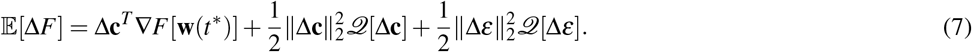

The requirement for assumption (6b) can be removed (see SI section 1.1). We can differentiate equation (7) in ‖Δ**c**‖_2_, to get:

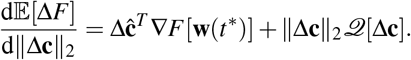

The root of this derivative gives a global minimum of the equation (7) in ‖Δ**c**‖_2_, as long as 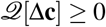 holds (justified in SI section 1.2). We get

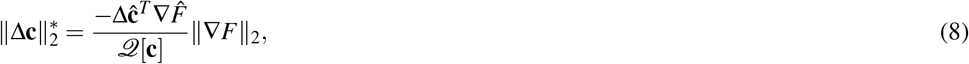

which defines the magnitude of systematic plasticity that minimises Δ*F*, and thus maximises task performance at time *t*\* + Δ*t*.

If the memory improved over the interval Δ*t*, then we would have Δ*F* < 0. However, we are interested in the special case of memory maintenance, where improvements from Δ**c** are cancelled out by decrements from Δ*ε*, and we have 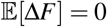. Substituting this into equation (7), we get

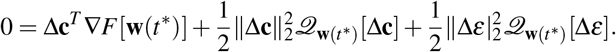

Next, we substitute in our optimal reconsolidation magnitude (equation (8)). This gives

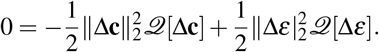

Therefore,

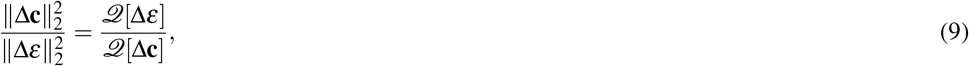

and moreover,

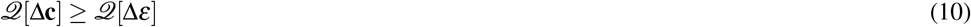

implies that the magnitude of synaptic fluctuation should outcompete that of systematic plasticity for optimal memory maintenance.

Note that an alternative derivation of equation (9) (see SI section 1.1) removes the need for assumption (6b). Equation (9) is valid when the numerator and denominator of the right hand side are both positive. The converse is unlikely in a partially trained network, and impossible in a highly trained network (see SI section 1.2).

Now let us provide more intuition for equation (9). Recall from Figure **??** that 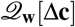 represents the relative upward curvature of *F*[**w**] in the direction Δ**c** (and the same for Δ*ε*). Therefore we can interpret (9) as follows:

Draw two unit-length lines on the loss function landscape, both starting at **w**(*t*\*), and in the directions 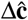 and 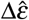 respectively. Measure the upward curvature of the loss function along both of these lines. If the curvature is greater (or equal) for line 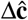, then optimal memory retention over Δ*t* requires 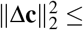 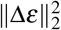.

Geometric intuition can give a glimpse into why equation (10) should generically hold, although we also demonstrate this mathematically (SI section 1.3). 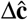 is a descent direction on the loss landscape, since it acts to improve memory quality (see equation (4)). Descent directions will generically have high upwards curvature in highly trained states (i.e. when *F*[**w**] is close to a minimum). In other words, a cross section of the loss landscape along such a descent direction will be U-shaped. For this to be false, *F* would have to consistently decrease along the cross section. However, the degree to which this decrease can occur is limited by the already low value of *F*[**w**]. On the other hand, arbitrary directions (such as Δ*ε*) may not act to decrease *F*, and thus may not have U-shaped cross sections.

We have intuitively justified why (10) should hold. This equation implies that systematic plasticity should generically be outcompeted by synaptic fluctuations, for optimal memory retention. We can justify the same assertion analytically by quantifying 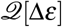 and 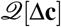.

First note that 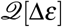 is task-independent. Thus, it should have no systematic relationship with directions of curvature at *F*[**w**]. In other words, it should project unbiasedly onto the different eigenvectors of ∇^2^*F*[**w**]. This assumption implies (see SI section 1.2)

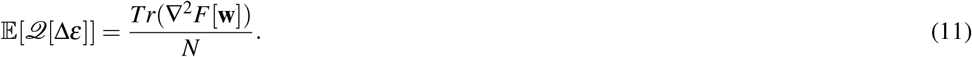

Note that the RHS of the above equation corresponds to the mean of the eigenvalues of ∇^2^*F*[**w**].

We now turn to quantifying 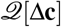. Regardless of the mechanism generating systematic plasticity, it must act to improve task performance. This constraint gave us equation (4): Δ**c** must anticorrelate with the gradient ∇*F*[**w**], and it can also benefit from information on ∇^2^*F*[**w**]. We can consider two extremal cases:

1. Δ**c** is computed with perfect access to ∇*F*[**w**], and (possibly) ∇^2^*F*[**w**].
2. The quantity of information on ∇*F*[**w**] available with which to compute Δ**c** tends towards zero.

In the SI (section 1.3), we show that synaptic fluctuations outcompete/equal systematic plasticity in both of these extremal cases, and intermediately by interpolation. We therefore suggest that in biological systems maintaining a memory through reconsolidation, the appropriate null hypothesis is that the magnitude of synaptic fluctuations outcompetes the magnitude of reconsolidation plasticity.

Another interesting phenomenon follows from the calculations of SI section 1.3, in the case that only information on ∇*F*[**w**] is avilable. If Δ**c** can perfectly access ∇*F*[**w**], then Δ**c** ∝ −∇*F*[**w**] optimally decreases *F*[**w**]. This corresponds to perfect implementation of backpropagation (i.e. gradient descent) by Δ**c**. In this case, (9) is smaller than one, and intrinsic fluctuations outcompete systematic plasticity. As we decrease the accuracy of backpropagation (i.e. corrupt access to ∇*F*[**w**] with task-independent noise), (9) increases towards one: the optimal magnitude of systematic plasticity increases. In other words, a less precise systematic plasticity mechanism has to do ‘more work’ to optimally counteract intrinsic fluctuations. This is demonstrated numerically in Figures 4 and 3.

**Figure 3.**
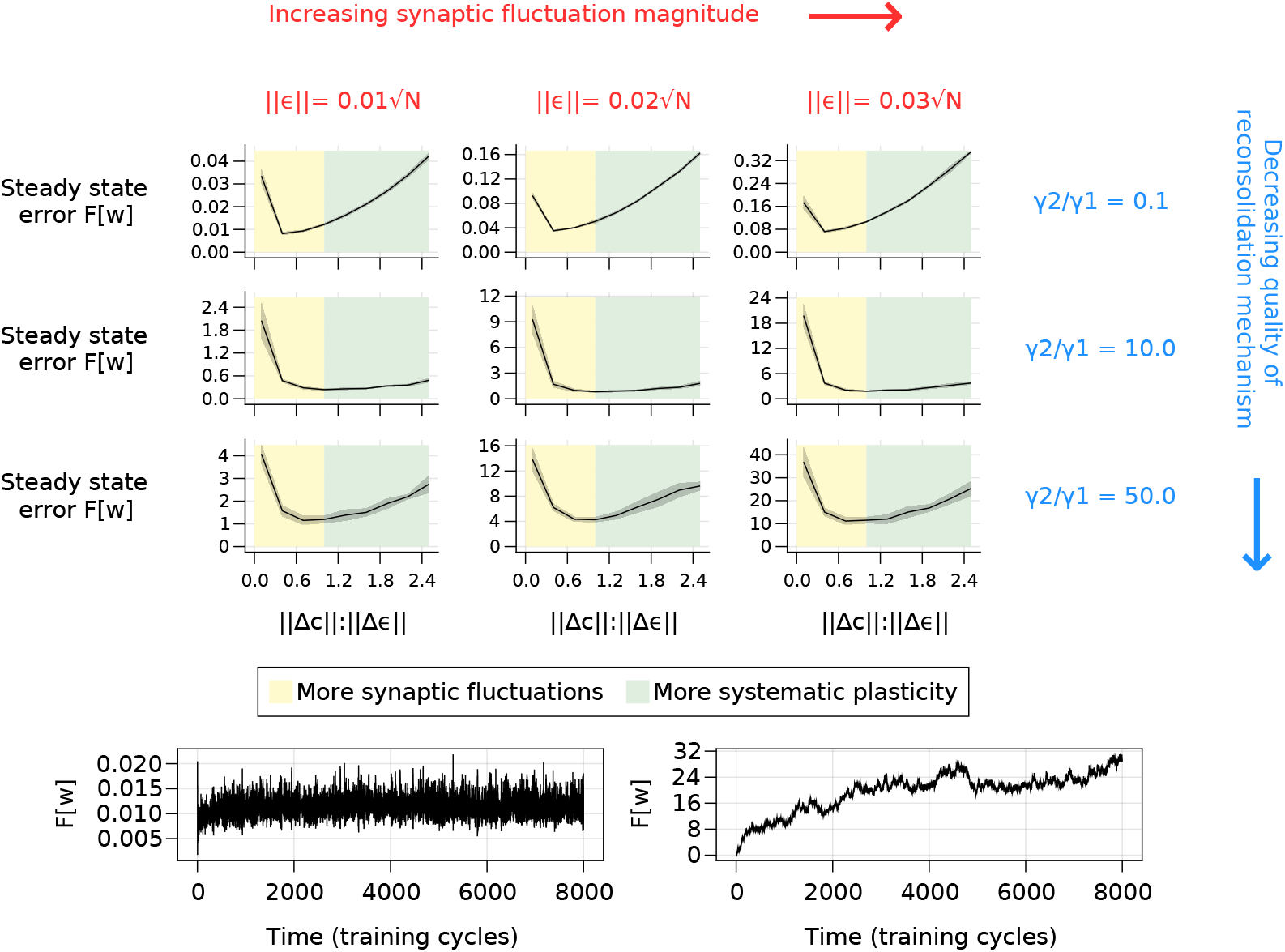
The relationship between steady state task performance and the ratio of systematic plasticity to intrinsic fluctuations in a nonlinear network. We consider multilayer perceptrons with 12 inputs, 10 outputs, and 20 neurons in the hidden layer. Each neuron has a sigmoidal nonlinearity. Weight dynamics are described in the ‘simulations’ section. Each subgraph in the top pane shows steady state error over 8 repeats, with standard deviation over the repeats shaded. The x-axis of each subgraph is equivalently 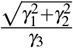. The bottom pane depicts sample trajectories of task error over time, for different choices of hyperparameters *γ*_1_, *γ*_2_, and *γ*_3_.

**Figure 4.**
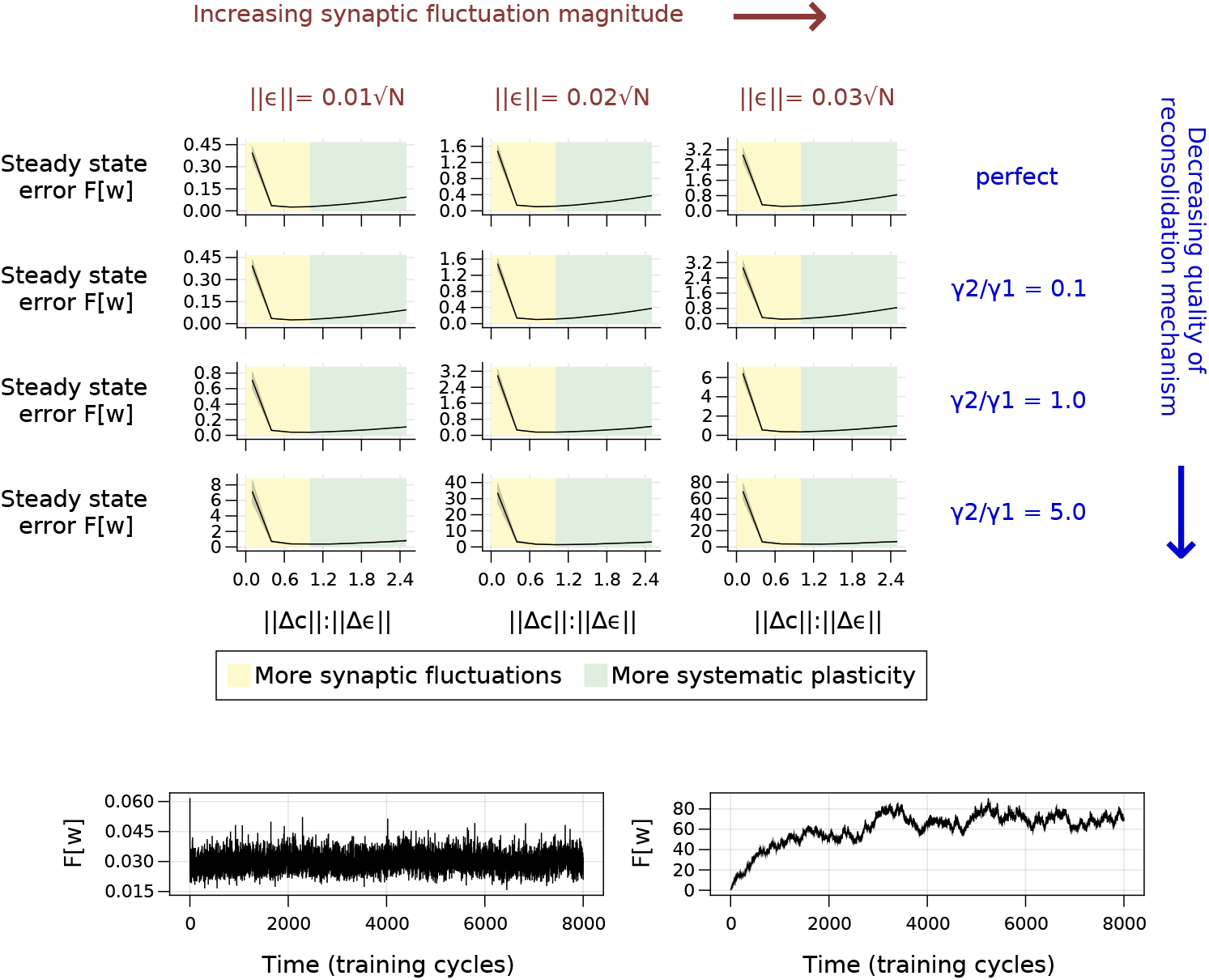
The relationship between steady state task performance and the ratio of systematic plasticity to intrinsic fluctuations in a linear network. We consider linear networks with output *y*(**w**, *u*) = *Wu*, where *W* ∈ ℝ^12×10^ is a matrix representation of the weight vector **w**. Weight dynamics are described in the ‘simulations’ section. We ran the dynamics for 8000 training cycles, to allow task performance to settle to a steady state. Each subgraph in the top pane shows steady state error over 8 repeats, with standard deviation over the repeats shaded. **Imperfect learning rules (columns 2-4)**: The x-axis of each subgraph is equivalently 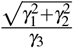. The bottom pane depicts sample trajectories of task error over time, for different choices of hyperparameters *γ*_1_, *γ*_2_, and *γ*_3_. **Perfect learning rule (top row)**: The systematic learning rule updates weights with the Newton step (Δ**c**_*t*_ ∝ (∇^2^*F*[**w**]_*t*_)^−1^*F*[**w**]_*t*_), which becomes (in the linear case): Δ**c**_*t*_ ∝ **w**\* − **w**. Overall weight dynamics are Δ**w**_*t*_ = Δ**c**_*t*_ + Δ*ε*_*t*_, as before.

We now verify our conclusions in simulations. We consider simple, feedforward, rate-based neural networks. We emphasise that the results do not depend on a particular choice of network model, learning rule, or task. Our aim is to verify the ratios of systematic to intrinsic plasticity that result in optimal steady-state performance on a particular learned task.

The task we provide our neural networks is to maintain their initial input-output behaviour over time as well as possible, even as individual weights fluctuate due to systematic and intrinsic plasticity. This models maintenance of a previously learned task. Initial network behaviour is set by randomly setting the initial neural network weights. We ‘save’ the initial network behaviour by keeping a fixed copy of the network at time zero, with fixed weights. We call this fixed network the ‘teacher’, and the adaptive network the ‘student’. Our learning problem then recreates the student-teacher framework described in e.g. Levin et al. (1990); Seung et al. (1992).

As before, we model weight change over a time interval *T* as

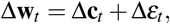

where the index *t* defines the number of previously elapsed intervals. We model Δ*ε*_*t*_, which represents integrated intrinsic fluctuations over the time interval, as a scaled, i.i.d, white-noise process at each synapse:

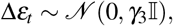

for some scaling factor *γ*_3_ > 0. In order to describe Δ**c**_*t*_ we first have to define an error function *F*[**w**] for the student network. We take

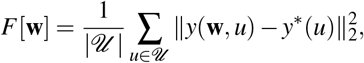

where 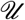 is a set of inputs with cardinality 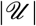, *y*\*(*u*) is the output of the teacher for input 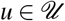, and *y*(**w**, *u*) is the output of the student, with weights **w**. We generated inputs 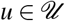 as i.i.d Gaussian vectors, and took 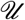 as a set of 1000 such vectors.

We model the systematic plasticity term Δ**c**_*t*_ as a noise-corrupted gradient descent term. Any such term must anticorrelate to some degree with the gradient (see equation (4) and surrounding discussion, as well as Raman et al. (2019)). So we take

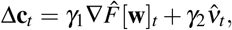

where 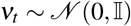 models imperfections in the approximation of the gradient, and *γ*_1_*, γ*_2_ > 0 are scaling parameters. By changing the ratio 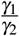, we can interpolate between high and low quality learning rules. Meanwhile 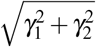 represents the overall magnitude of systematic plasticity in the time interval, which can be compared with *γ*_3_, the magnitude of intrinsic fluctuations.

We ran simulations of a sigmoidally nonlinear multilayer perceptron with a single hidden layer, tuning the vector *γ* of hyperparameters to investigate different qualities of learning rule, and different ratios of systematic to intrinsic fluctuations (see Figure 3). Under all conditions, optimal steady state performance was achieved when the magnitude of systematic plasticity was less or equal to the level of intrinsic fluctuations, corroborating analytical calculations.

## DISCUSSION

A long-standing question in neuroscience is how neural circuits optimally maintain memory of a learned task while being buffeted by synaptic fluctuations from noise and other task-independent processes (Fusi et al., 2005). There are several hypotheses that offer potential answers, none of which is mutually exclusive. One possibility is that fluctuations only occur in a subset of volatile connections that are relatively unimportant for learned behaviours Moczulska et al. (2013); Chambers and Rumpel (2017); Kasai et al. (2003). Following this line of thought, circuit models have been proposed that only require stability in a subset of synapses for stable function Clopath et al. (2017); Mongillo et al. (2018); Susman et al. (2018). Another hypothesis is that any memory degradation due to fluctuations is counteracted by restorative, systematic plasticity processes that allow circuits to continually ‘relearn’ stored associations. The source of information for the systematic plasticity term could come from an external reinforcement signal Kappel et al. (2018), from interactions with other circuits Acker et al. (2018), or spontaneous, network-level reactivation events Fauth and van Rossum (2019). A final possibility is that we rarely observe behavioural states in which an animal is not learning, and that unobserved behavioural changes account for apparent fluctuations in brain connectivity in any given experiment.

Our work does not argue exclusively for or against any of these three broad hypotheses. Rather, we extracted logical consequences from assuming that all hypotheses are viable to some extent. An important caveat to our work is that we do make specific assumptions whose validity depends on the state of current knowledge, and might vary depending on the organism or brain area in question. Most crucially, we assumed that not all fluctuations in synaptic strength are explained by behavioural adaptation and learning. To the extent that this is true, the residual fluctuations must come from ongoing, endogenous processes and irreducible noise that continually perturb synaptic strengths. Our analysis then proceeded by assuming that learned information does not decay appreciably over time, which requires degradation from these task-independent processes to be counteracted by systematic plasticity mechanisms.

Several predictions follow immediately from our analysis. Foremost among these, we predict that for optimal retention of circuit function and learned behaviour, the systematic plasticity contribution should not outcompete task-independent fluctuations. This prediction is somewhat unsettling yet it is borne out across a number of experimental studies (Nagaoka et al., 2016; Quinn et al., 2019; Yasumatsu et al., 2008; Minerbi et al., 2009; Dvorkin and Ziv, 2016). It is intuitively clear that memories should degrade when systematic plasticity is far weaker than noise. It is far less intuitive that maintenance of learned behaviours will also suffer if a learning rule outcompetes task-independent fluctuations at steady state.

By parameterising all possible qualities of systematic plasticity - from precise to highly inaccurate - we also show that a larger task-independent component of ongoing synaptic change predicts a more accurate systematic plasticity mechanism. In other words, sophisticated learning rules need to do less work to overcome the damage done by task-independent synaptic fluctuations. Experimental estimates (see Table 1) suggest task-independent fluctuations often outcompete systematic changes in synaptic strength. Our theory implies that this is consistent with relatively precise learning rules in biological synapses that can approximate the gradient of task error relatively well.

We must qualify the meaning of ‘relatively precise’ in this conclusion. We conceptualised memory quality as a landscape whose height denotes error. We noted that any systematic plasticity mechanism should act to increase memory quality, and therefore change the weights in a downhill direction on the landscape. Therefore it must have at least some local information on the slope (gradient) and possibly curvature (hessian) of the landscape. We parameterised precision by the quality of access to these quantities. The assertion of the previous paragraph assumed no access to the curvature, as is consistent with biologically plausible learning rules we have seen in the literature (e.g. Williams (1992); Mazzoni et al. (1991); Seung (2003); Lillicrap et al. (2016); Sussillo and Abbott (2009)).

If it were the case that biological learning rules could perfectly access both the gradient and second derivative of the landscape, then the optimal contributions of systematic plasticity and intrinsic fluctuations would in fact be equal (SI section 1.3.2). This would correspond to the systematic plasticity mechanism directly undoing any synaptic changes induced by task-independent fluctuations. We suggest that this is not biologically realistic. First, it is widely believed that even accurate gradient-based learning rules are biologically implausible. Second, there is widespread evidence that neural circuit reconfiguration occurs in the absence of learning and that many synapses have finite lifetimes, indicating that synapses do not continually revert to some learned state.

There are important caveats to how our results should be interpreted in light of existing experimental data. It is technically difficult to experimentally isolate systematic and random components of synaptic change. Approaches often rely on reduced preparations where ‘learning’ and ‘behaviour’ have no direct relationship to neural circuit dynamics. On the other hand, in *in vivo* studies it is extraordinarily difficult to accurately measure synaptic changes and to control for confounding changes in behaviour or physiology. We may therefore simply take these measurements as the best available data and assume that *at least some* ongoing synaptic dynamics are noise-driven.

Thus, while our results offer a surprising agreement with a number of experimental observations, we believe it is important to further replicate measurements of synaptic turnover and synaptic modification in a variety of settings, both *in vivo* and *in vitro*. To this end, we hope our results provide an impetus for this difficult experimental work, because it offers a principled framework for understanding the surprising volatility of connections in neural circuits.

## ACKNOWLEDGMENTS

This work was supported by ERC grant StG 2016 716643 FLEXNEURO.

## 1 SUPPLEMENTAL METHODS

### 1.1 Alternative derivation of equation (9)

We provide an alternative derivation of equation (9) that removes the need for assumption (6b). We did not put this main derivation in the main text as we perceive it to have less clarity.

The derivation proceeds identically to that given in the main text until equation (5). We can then use (6a) to simplify equation (5). We get

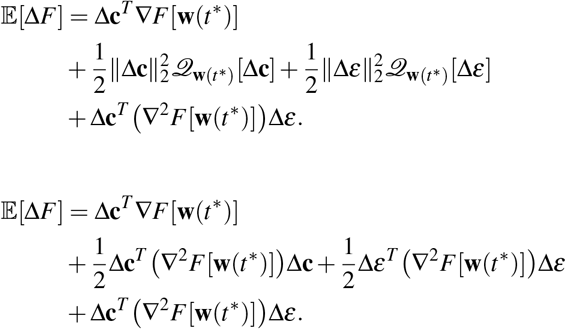

Recall that expectation is taken over an unknown probability distribution from which Δ*ε* is drawn, which satisfies equation (6a).

We then assume that we are in a phase of stable memory retention, so that 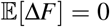. Now if the magnitude of systematic plasticity ‖Δ**c**‖_2_ is tuned to minimise steady state error *F*, then any change to ‖Δ**c**‖_2_ will result in an increase in 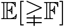. So 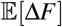 is locally minimal in ‖Δ**c**‖_2_. This implies

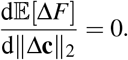

We also claim that local minimality implies

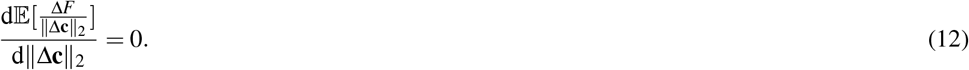

Why? 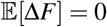 implies that 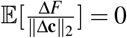. If a small change to ‖Δ**c**‖_2_ results in 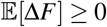, then it also results in 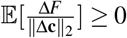, since ‖*dC*‖_2_ is non-negative.

Expanding the LHS of equation (12), we get

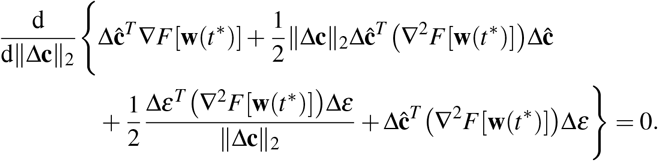

Differentiating, we get

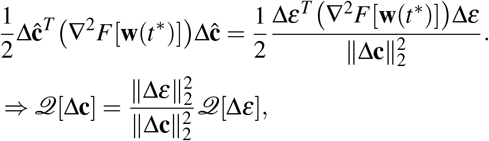

from which (9) follows.

### 1.2 Positivity of the numerator and denominator in equation (9)

Equation (9) of the main text asserts that

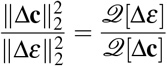

holds as long as both the numerator and denominator of the RHS are positive. Here we describe sufficient conditions for positivity.

The inequality ∇^2^*F*[**w**] ⪰ 0 must hold in some neighbourhood of any minimum **w**\*. Recall that we referred to such a neighbourhood as a highly trained state of the network in the main text. In such a state, our assertion follows immediately, as 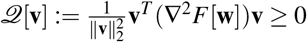, for any vector **v**. Therefore 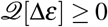 and 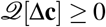.

We now consider a partially trained network state, which we defined in the main text as any **w** satisfying *Tr*(∇^2^*F*) ≥ 0. Note that

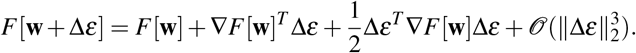

We assumed in the main text (equation (1a)), that Δ*ε* is uncorrelated with the gradient ∇*F*[**w**] in expectation, since Δ*ε* is realised by memory-independent processes. Similarly we can assume that Δ*ε* is unbiased in how it projects onto the eigenvectors of ∇^2^*F*[**w**]. In other words,

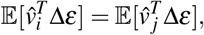

for any normalised eigenvectors *v*_*i*_, *v* _*j*_ of ∇^2^*F*[**w**]. In expectation, we can therefore simplify to

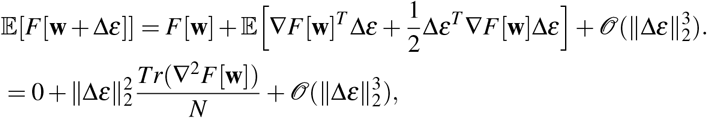

where *N* is the dimensionality of the vector **w**. So a partially trained network is one for which small, memory-independent weight fluctuations (such as Δ*ε*, or white noise) are expected to decrease memory quality.

Now recall that 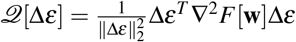. So we have

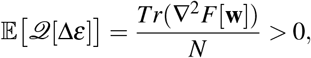

where the positivity constraint comes from being in a partially trained network.

We now consider why 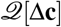 should be generically positive in a partially trained network. Suppose 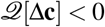 holds. We can rewrite this as Δ**c**^*T*^ ∇^2^*F*[**w**]Δ**c** ≤ 0. In this case, maintaining the same systematic plasticity direction Δ**c** over the time interval [*t*\* + Δ*t, t*\* + 2Δ*t*] would result in increased improvement in loss, as

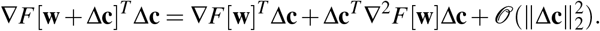

Effectively, memory improvement due to systematic plasticity Δ**c** would be in an ‘accelerating’ direction, and maintaining the same direction Δ**c** of systematic plasticity would lead to ever faster learning. However, by assumption, we are in a regime of steady state task performance, where

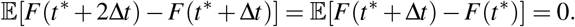

### 1.3 Optimal plasticity ratios in specific learning rules

#### 1.3.1 Noise-free learning rules (first-order)

Let us first consider the case where Δ**c** can be computed with perfect access to the gradient ∇*F*[**w**], but without access to ∇^2^*F*[**w**]. Such a Δ**c** is known as a first-order learning rule, as it has access only to the first derivative of *F* Polyak (1987). Imperfect access is considered subsequently. In this case, the optimal direction of systematic plasticity is

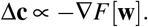

In other words, Δ**c** would implement perfect gradient descent on *F*[**w**]. The condition (10) for synaptic fluctuations to outcompete reconsolidation plasticity evaluates to

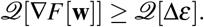

To what extent can we quantify 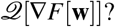 First let us relate the gradient and Hessian of *F*[**w**]. Let **w**\* be an optimal state of the network (i.e. one where *F* is minimised). Let us parameterise the straight line connecting **w** with **w**\*:

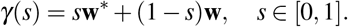

Then

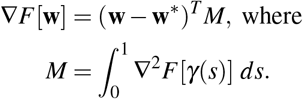

This gives

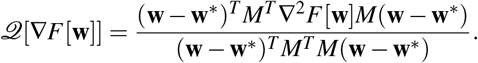

First let us rewrite

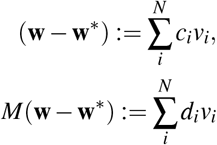

where (*λ*_*i*_, *v*_*i*_) is the *i*^*th*^ eigenvalue/eigenvector pair of ∇^2^*F* (sorted in ascending order of *λ*_*i*_), and *c*_*i*_, *d*_*i*_ are some scalar weights. Now

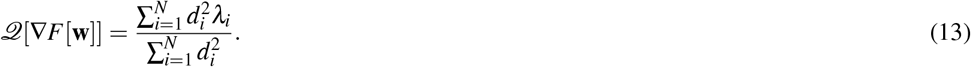

The value of 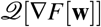 now depends upon the distribution of mass of the sequence {*d*_*i*_}. If later elements of the sequence are larger (i.e. *M*(**w**= **w**\*) projects more highly onto eigenvectors of ∇^2^*F*[**w**]) with large eigenvalue), then 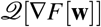 becomes larger, and the optimal magnitude of reconsolidation plasticity decreases, relative to the magnitude of synaptic fluctuations. The opposite is true if earlier elements of the sequence are larger.

Guaranteed bounds on the value of equation (13) are vacuous. If we do not restrict *M*, then we can tailor the sequence {*d*_*i*_} as we like, and we end up with 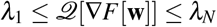. However, pragmatic bounds are much tighter. Let us now consider two plausibly extremal cases.

First consider the simplest case of a network that linearly transforms its outputs, and which has a quadratic loss function *F*[**w**]. In this case ∇^2^*F* is a constant, (independent of **w**), positive-semidefinite matrix, and *M* = ∇^2^*F*. This means that

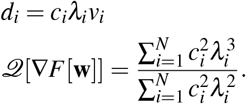

Condition (10) then becomes

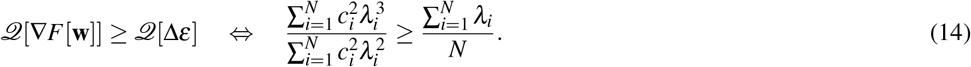

A conservative sufficient condition for (14), using Chebyshev’s summation inequality, is that

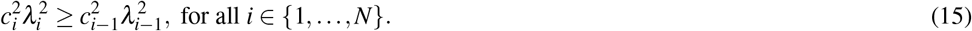

Under what conditions would a plausible reconsolidation mechanism choose to ‘outcompete’ synaptic fluctuations, in this linear example? For 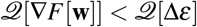 to even hold, (15) would have to be broken, and significantly so due to conservatism in the inequality. In other words, **w** − **w**\* must project quite biasedly onto the eigenvectors of ∇^2^*F* with smaller-than-average eigenvalue. If the discrepancy between **w** and **w**\* were caused by fluctuations (which are independent of ∇^2^*F*), then this would not be the case, in expectation. Even if this were the case, the reconsolidation mechanism would have to know about the described bias. This requires knowledge of both **w**\* and ∇^2^*F*, and is thus implausible.

Now let us consider the case of a generic nonlinear network. At one extreme, if ‖**w** − **w**\*‖_2_ is small, then *M* ≈ ∇^2^*F*[**w**], and the discussion of the linear case is valid. This corresponds to the case where steady state error is close to the minimum achievable by the network. As ‖**w** − **w**\*‖_2_ increases (i.e. steady state error gets worse), the correspondence between *M* and ∇^2^*F*[**w**] will likely decrease. Thus the optimal magnitude of reconsolidation plasticity, relative to the level of synaptic fluctuations, will rise.

We could consider another ‘extreme’ case in which *M* and ∇^2^*F*[**w**] were completely independent of each other. In this case,

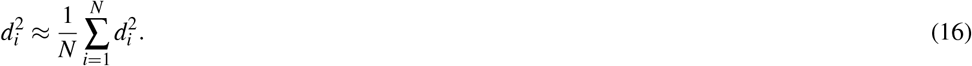

In other words, the projection of *M*(**w** − **w**\*) onto the different eigenvectors of ∇^2^*F*[**w**] is approximately even. Using (13), this gives

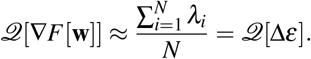

In summary, we have two plausible extremes. One occurs where *M* = ∇^2^*F*[**w**], and another occurs where *M* is completely independent of ∇^2^*F*[**w**]. In either case, 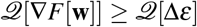, and so the magnitude of synaptic fluctuations should optimally outcompete/equal the magnitude of reconsolidation plasticity. Of course, there might be particular values of **w** where the correspondence between *M* and ∇^2^*F*[**w**] is ‘worse’ than chance. In other words, eigenvectors of *M* with large eigenvalue preferentially project onto eigenvectors of ∇^2^*F*[**w**] with small eigenvalue. In such cases, we would have 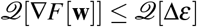. However, we find it implausible that a reconsolidation mechanism would be able to gain sufficient information on *M* to determine this at particular points in time, and thereby increase its plasticity magnitude.

#### 1.3.2 Noise-free learning rules (second order)

Let us now suppose that Δ**c** can be computed with perfect access to both ∇*F*[**w**] and ∇^2^*F*[**w**]. In this case the reconsolidation mechanism would optimally apply plasticity in the direction of the Newton step: we would have

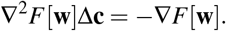

Note that the Newton step is often conceptualised as a weighted form of gradient descent, where movement in weight space is biased towards direction of lower curvature. Thus we would expect 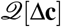 to be smaller, and the optimal proportion of reconsolidation plasticity to be larger. This is indeed the case. For mathematical tractability, we will restrict our discussion to the case in which ∇^2^*F*[**w**] ≻ 0, and *M* ≻ 0. This would hold if *F*[**w**] were convex, or if **w** were sufficiently close to a local minimum **w**\*. In this case we can rewrite

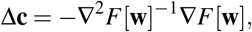

which gives

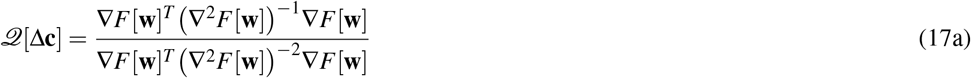

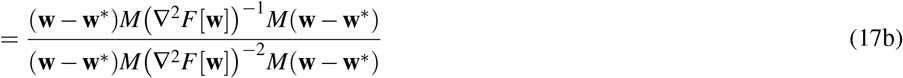

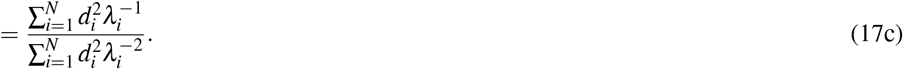

Once again, we first consider the case of a linear network with quadratic loss function, and hence with constant Hessian ∇^2^*F*. This gives *M* = ∇^2^*F*, and

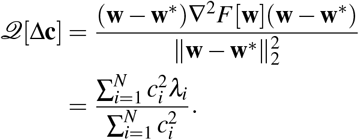

We again assume that the reconsolidation mechanism does not have knowledge of the relative projections of **w** − **w**\* onto the different eigenvectors of ∇^2^*F*, which requires knowledge of **w**\*. Without such information, we can use an analogous argument to that preceding (16) to argue that the approximation 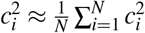 is reasonable. This gives 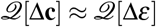.

Note that the Newton step, in the linear-quadratic case just considered, corresponds to a direction **w**\* − **w**, i.e. a direct path to a local minimum. So we could consider a systematic plasticity mechanism implementing the Newton step as one directly undoing synaptic changes caused by the intrinsic flucatuations Δ*ε*.

We now consider the case of a nonlinear network. As before, if ‖**w** − **w**\*‖_2_ is small, then we have *M* ≈ ∇^2^*F*[**w**], and the arguments of the linear network hold. As ‖**w** − **w**\*‖_2_ increases, the correspondence between *M* and ∇^2^*F* will decrease. We again consider the plausible extreme where *M* is completely uncorrelated with ∇^2^*F*[**w**], and so the approximation (16) holds. In this case, equation (17c) can be simplified to give

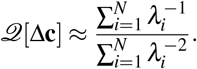

We assumed that ∇^2^*F*[**w**] ≻ 0. Therefore, all eigenvalues are positive. This allows us to use Chebyshev’s summation inequality to arrive at

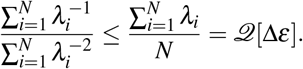

So as ‖**w** − **w**\*‖_2_ increases, the magnitude of reconsolidation plasticity will optimally outcompete that of synaptic fluctuations.

#### 1.3.3 Imperfect learning rules

The previous section applied in the implausible case where a reconsolidation mechanism had perfect access to ∇*F*[**w**] and/or ∇^2^*F*[**w**]. Recall, from the discussion surrounding equation (4), that at least some information on ∇*F*[**w**] is required. What if Δ**c** contains a mean-zero noise term, corresponding to imperfect access to these quantities? We will now show how such noise pushes 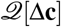 towards equality with 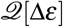, and thus pushes the optimal magnitude of reconsolidation plasticity towards the magnitude of synaptic fluctuations. Let us use the model

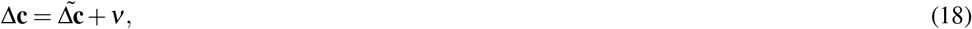

where *ν* is some mean-zero term, and 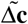 is the ideal output of the reconsolidation mechanism, assuming perfect access to the derivatives of *F*[**w**]. Here *ν* represents the portion of systematic plasticity attributable to systematic error in the algorithm, due to imperfect information on *F*[**w**]. This could arise due to imperfect sensory information or limited communication between synapses. We can therefore assume, as for Δ*ε*, that it does not contain information on ∇^2^*F*[**w**]. We therefore get

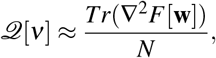

analogously to (11). Now the operator 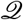 satisfies

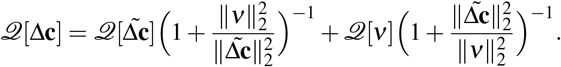

So depending upon the relative magnitudes of 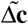 and *ν*, 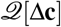 interpolates between 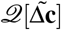 and 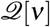. In particular, as the crudeness of the learning rule (i.e. the ratio 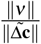) grows, 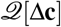 approaches equality (from below) with 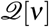, and thus 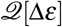, completing our argument.

